# Dynamics-function relationship of N-terminal acetyltransferases: the β6β7 loop modulates substrate accessibility to the catalytic site

**DOI:** 10.1101/502799

**Authors:** Angèle Abboud, Pierre Bédoucha, Jan Byška, Thomas Arnesen, Nathalie Reuter

**Affiliations:** Department of Informatics, University of Bergen, Bergen, Norway; Computational Biology Unit, Department of Informatics, University of Bergen, Bergen, Norway; Department of Biological Sciences, University of Bergen, Bergen, Norway; Department of Biomedicine, University of Bergen, Bergen, Norway; Department of Surgery, Haukeland University Hospital, Bergen, Norway; Department of Chemistry, University of Bergen, Bergen, Norway

**Keywords:** Acetylation, N-terminal acetyltransferases, protein dynamics, normal modes analysis, ligand specificity

## Abstract

N-terminal acetyltransferases (NATs) are enzymes catalysing the transfer of the acetyl from Ac-CoA to the N-terminus of proteins, one of the most common protein modifications. Unlike NATs, lysine acetyltransferases (KATs) transfer an acetyl onto the amine group of internal lysines. To date, not much is known on the exclusive substrate specificity of NATs towards protein N-termini. All the NATs and some KATs share a common fold called GNAT. The main difference between NATs and KATs is an extra hairpin loop found only in NATs called β6β7 loop. It covers the active site as a lid. The hypothesized role of the loop is that of a barrier restricting the access to the catalytic site and preventing acetylation of internal lysines. We investigated the dynamics-function relationships of all available structures of NATs covering the three domains of life. Using elastic network models and normal mode analysis, we found a common dynamics pattern conserved through the GNAT fold; a rigid V-shaped groove, formed by the β4 and β5 strands and three relatively more dynamic loops α1α2, β3β4 and β6β7. We identified two independent dynamical domains in the GNAT fold, which is split at the β5 strand. We characterized the β6β7 hairpin loop slow dynamics and show that its movements are able to significantly widen the mouth of the ligand binding site thereby influencing its size and shape. Taken together our results show that NATs may have access to a broader ligand specificity range than anticipated.

**Author summary:** N-terminal acetylation concerns 80% of eukaryotic proteins and is achieved by enzymes called the N-terminal acetyltransferases (NATs). They belong to the large family of acetyltransferases and adopt the GNAT fold. Interestingly most lysine acetyltransferases (KATs), which acetylate specifically internal lysines, share the same fold. Rationale for the ligand recognition by the GNAT enzymes remains unclear. Proteins are dynamic entities that utilize their structural flexibility to carry out functions in living cells. By studying the dynamics throughout the entire NATs family, we found that the slow dynamics of the fold is strongly conserved. We also revealed the mobility of the active site lid, namely the β-hairpin loop β6β7, which is one of the main structural differences between the NATs and the KATs. The size and shape of the ligand binding site depend on movements of that β-hairpin loop. We suggest that in attempts of mapping NATs specificity or ligand design the fold flexibility should be taken into consideration.

## Introduction

Acetyltransferases are enzymes catalysing the transfer of an acetyl group from the co-factor acetyl-coenzyme A (Ac-CoA) to a substrate. Among them, lysine acetyltransferases (KATs) and Να-terminal acetyltransferases (NATs) perform protein acetylation to either lysine side chains or N-termini of polypeptide chains, respectively. NATs acetylate 80 to 90% of the proteins of the human proteome [1] and N-terminal acetylation has been shown to play a role in various biological processes from protein folding to gene regulation [2]. Dysregulation or mutations of NATs have been linked to several diseases including tumour development [2–5] and initiatives are already undertaken to develop inhibitors targeting the relevant NATs [6].

Most acetyltransferases share the GNAT fold (Gcn5-related N-acetyltransferases) [7]. It consists of a three-layered αβα sandwich containing seven β-sheets and four α-helices (Fig. 1). The GNAT fold displays two features that are conserved through most of the KATs and NATs and are related to the transfer of an acetyl to an amino group. The first is a conserved sequence motif essential for Ac-CoA binding (Q/RxxGxG/A) and located on the turn between strand β4 and helix α3 [7–9]. The β4 strand together with β4-α3 and α4 form much of the Ac-CoA binding site. The second salient feature of the GNAT fold is the V-shaped configuration of the two parallel strands β4 and β5, forming a groove where the extremities of the Ac-CoA and of the substrate peptide meet, positioning the acetyl group and the amino group close enough for the catalytic reaction to occur [7].

**Fig 1.**
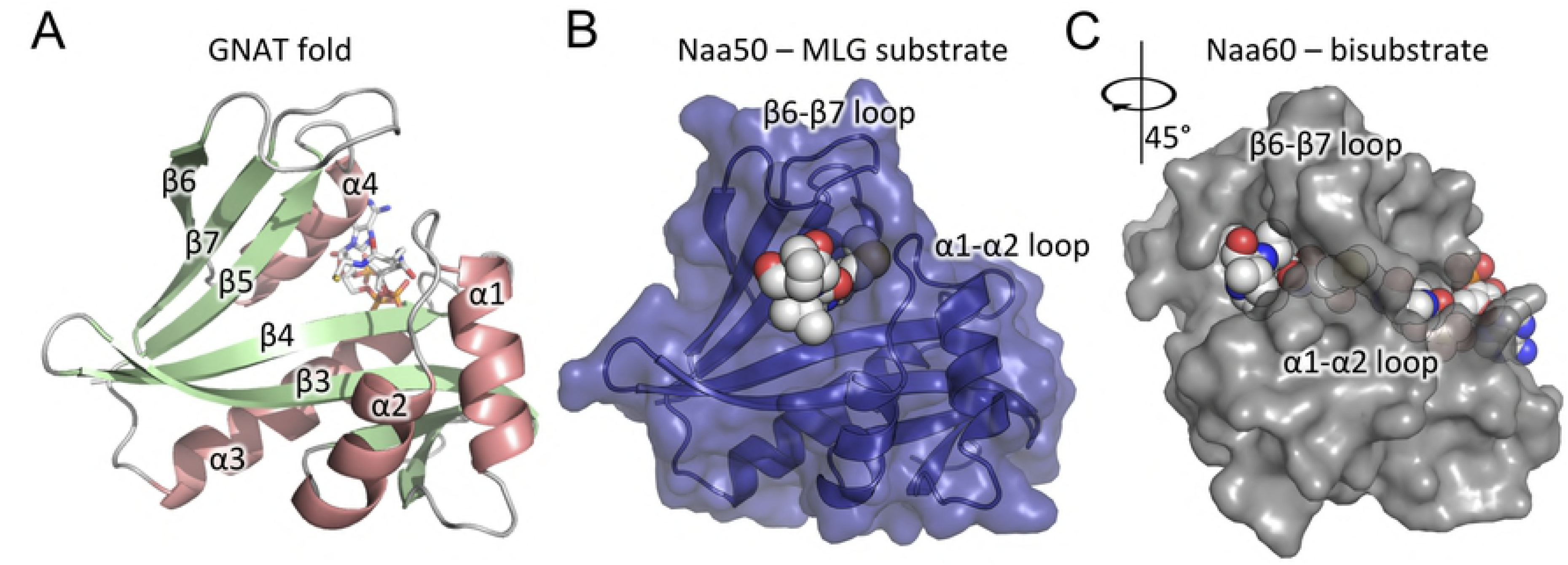
GNAT fold and substrate binding sites in NAT catalytic domains. (A) Cartoon representation of the GNAT fold of human Naa50 (PDB ID: 3TFY). It consists of 6 β-sheets (green) and 4 α-helices (salmon pink) organised in the following topology: B1-H1-H2-B2-B3-B4-H3-B5-H4-B6. The Ac-CoA represented in sticks with the backbone in grey sits between helix α4 and the β4α3 loop. (B) Human Naa50 (cartoons and blue solvent-accessible surface) bound to substrate Met-Leu-Gly (van der Waals spheres) (PDB ID: **3TFY**). (C) Naa60 (grey solvent-accessible surface) bound to a bisubstrate CoA-Ac-Met-Lys-Ala-Val. Loops β6β7 and α1α2 are labelled in (B) and (C).

Substrate specificity varies drastically within the NATs, offering a palette of enzymes able to target a large spectrum of N-terminal sequences. As of now, eight NATs (NatA-NatH) have been identified in eukaryotes [2,10,11], three in prokaryotes (RimI, RimJ and RimL) [12,13] and one in archaea (referred here as ArNat) [14]. NATs are classified based on their composition and substrates. Comparison of structures of NATs resolved by X-ray crystallography and KATs revealed the presence of a seventh β-strand at the C-terminus as the main conserved characteristic of the NATs [15–17]. The extra β-hairpin β6β7, the α1α2 loop and the helix α2 contain amino acids forming the boundary of the binding pocket, which we hereafter refer to as the *mouth* of the binding pocket. NAT substrates usually position two to three residues in the peptide binding site [15–18]. Two conserved tyrosines located on the β6β7 loop and another one on the α1α2 loop have been shown to interact with the substrate backbone via hydrogen bonds [19]. Both loops cover the groove containing the catalytic site (Fig. 1B) to which substrate peptide and Ac-CoA bind from opposite sides. The loops have been proposed to prevent the access of internal lysine to the catalytic site and as a consequence prevent their acetylation by NATs [15,20]. However, there have been reports of lysine acetylations by NATs [18,21–27]. Moreover, NATs can be inhibited by so-called bisubstrate inhibitors consisting of a short polypeptide covalently bound to the Ac-CoA [6,17,28,29]. The X-ray structure of the human NatF bound to bisubstrate CoA-Ac-MKAV_7_ shows that the inhibitor is placed in the Ac-CoA and substrate binding site with the β6β7 hairpin loop hanging over the top of it [28] (Fig. 1C). This structure raises the question of how the bisubstrate accesses both the ligand and Ac-CoA binding sites and as a consequence suggests the mouth of the active site needs to be able to open up.

Using relatively short 10 nanoseconds-long molecular dynamics simulations of the catalytic domains of two human NATs, we earlier showed for hNaa50 that (i) helix α2 undergoes flexibility changes upon ligand binding and (ii) the β6β7 hairpin loop was the most flexible region of the protein [19]. Likewise, we observed the β6β7 loop of the human Naa10 to be flexible [30] in 100 ns-long molecular simulations. The time-scale of these simulations did not allow us to investigate conformational changes happening on longer time-scales, such as displacements of loops. Enzyme dynamics is known to be important for their function. Notably, catalytic residues are reported to be placed at rigid positions of the protein structure, while amino acids involved in substrate binding are likely to have a higher flexibility [31–33]. Furthermore, functionally relevant flexibility is conserved between enzymes sharing the same fold. Normal mode analysis (NMA) using elastic network models (ENM) is an efficient computational method that has proven reliable to characterize the flexibility properties intrinsic to protein structures [34–40]. It has also been successfully used to conduct comparative analyses of multiple protein structures [31,34].

In this study, we characterize the dynamics of all available structures of NATs and compare them to uncover dynamics patterns intrinsic to the GNAT fold and relevant for function.

## Results

Our dataset consists of 34 structures of 15 distinct proteins listed in Table 1 and is representing ten types of NATs defined according to their composition and substrates (S1 and S2 Tables). The dataset spans the three domains of Life, where six out of eight NATs stem from eukaryotes (NatA, NatB, NatD, NatE, NatF, NatH), three from bacteria (RimI, RimJ and RimL) and one from archaea (ArNat). In what follows, the ten types of NATs found in our dataset are referred to by the name of the catalytic subunit of each NAT complex (Naa10, Naa20, etc … reported in Table 1 in the column titled “Group”).

**Table 1.**
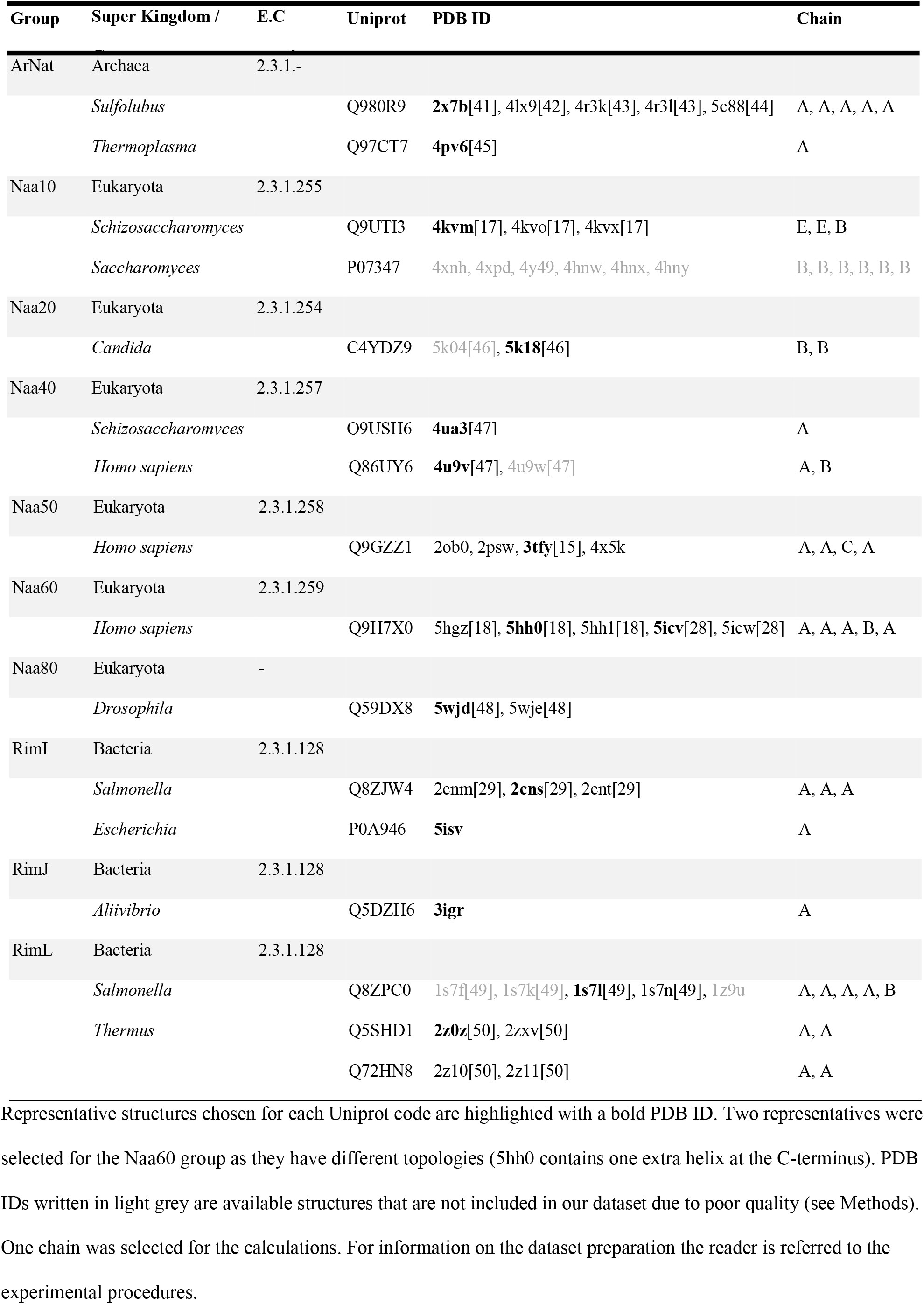
N-terminal acetyltransferases (NATs) found in PROSITE, Uniprot and PDB databases.

### (1) The NATs’ functional diversity builds upon the GNAT fold and is fine-tuned by accessory structural elements

We aligned all structures in the dataset using the multiple structure alignment tool MUSTANG [51] (see Experimental procedures). The structure of Naa50 (PDB ID: **3TFY**) was used as reference. The alignment led to 119 C-alpha atoms’ positions conserved (Fig. 2A). While sequence similarity between pairs of NATs is relatively low (23% identity on average), the secondary structure elements of the GNAT fold align well (Fig. 2B). The region between the end of strand β4 and helix α4 is the most conserved sequence-wise. It contains several of the amino acids involved in catalysis and located on strands β4 and β5, as well as the Ac-CoA binding motif (R/QxxGxG/A on β4-α3) (Fig. 2B). Noticeably, position 213 of the alignment (Asn114 in Naa50) is an asparagine conserved through all the 14 NATs, except in Naa80 where it is replaced by an aspartate (Asp 127) (Fig. 2B). This residue sits close to the oxygen of the Acetyl-CoA pantothenic acid for all structures [43]. Structural differences between NATs are restricted primarily to the N- and C-terminal regions, before helix α2 and after strand β6, respectively. The differences stem either from the longer elements of the GNAT domain or from additional accessories. The latter are secondary structure elements that are not part of the GNAT fold such as the N-terminal helix in Naa40, referred to as helix α0 or the sixty-one amino acid-long extra C-terminal helical segment in Naa60. Only the first thirty amino acids are resolved in the X-ray structure (α5 on Fig. 2C) but two helices are predicted from sequence analysis [52].

**Fig 2.**
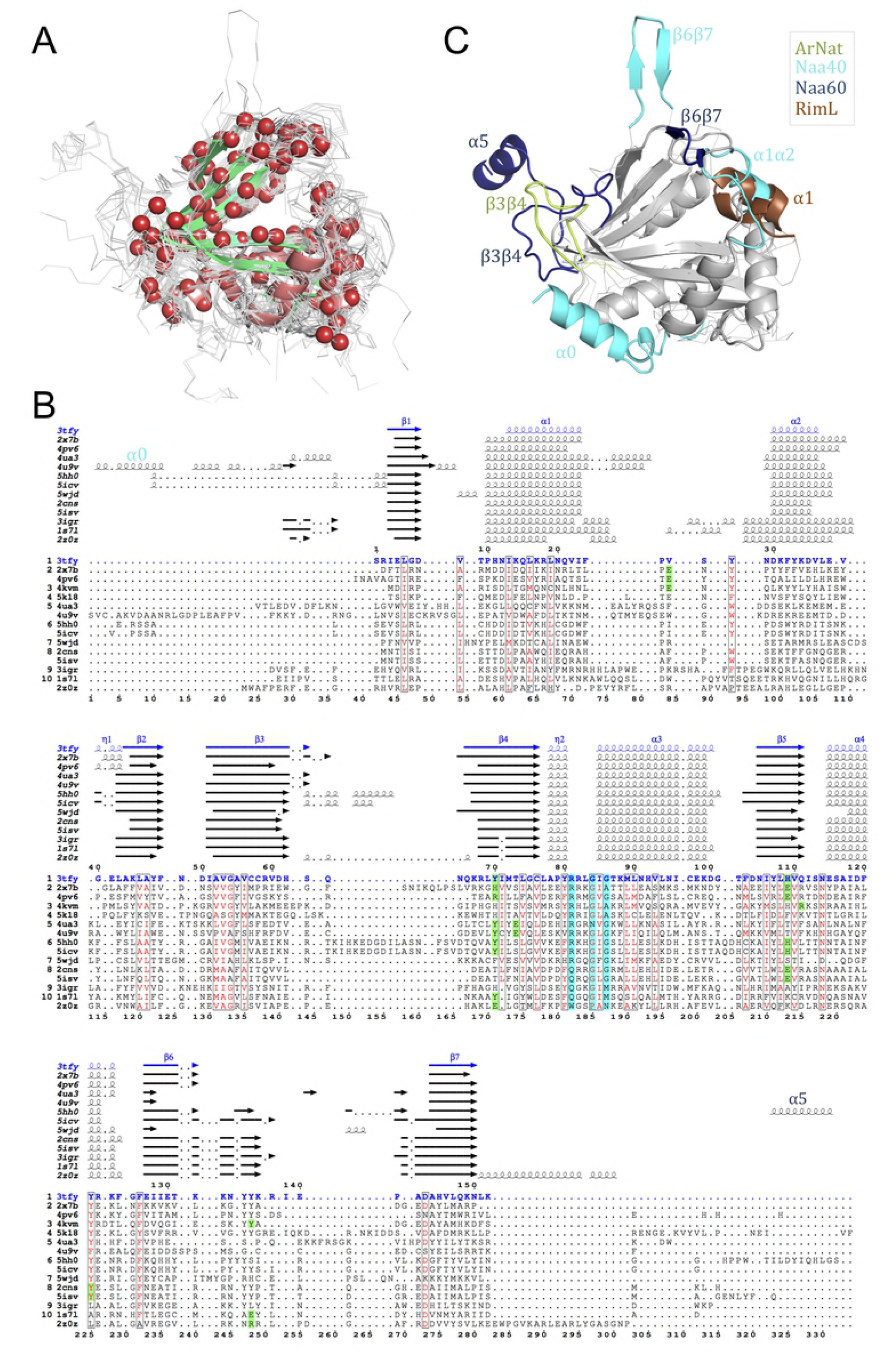
Structural alignment of Naa representatives. (A) MUSTANG structural alignment of all NAT structures listed in Table 1. The backbone of each structure is represented with lines except for that of the reference structure Naa50 (PDB ID: **3TFY**), which is represented with cartoons. The red beads represent the 116 aligned C-alpha atoms. (B) Multiple sequence alignment resulting from the structural alignment. Naa50 sequence is written with blue fonts, Ac-CoA-binding motifs are highlighted with cyan boxes and residues involved in the catalytic activity with green boxes. Sequences are labelled with the PDB ID from which their secondary structure elements are retrieved. All are shown above the sequence alignment except for *S. cerevisiae* Naa10 due to poor secondary structure annotation (PDB ID: **4Y49**). The image results from the use of ESPript [53]. (C) Cartoon representation of the shared GNAT fold (in grey) and structural variations: helix α0 in Naa40, helix α5 in Naa60, long β6β7 loops in Naa40 and Naa60, long α1α2 loops in Naa40 and RimL, long β3β4 loops in ArNats and Naa60.

We quantified the structural similarity between Naas by calculating pairwise root mean square deviations (RMSD) between the thirty-four structures of the dataset. The clustered values presented on a heatmap reveal two main groups (Fig. 3). As expected, RMSD values between structures belonging to the same group are below 1Å, which is in agreement with the fact that structures within a group are orthologues and/or structures of the same protein but in different forms (e.g. apo vs. holo) (S1 Table). To evaluate the influence of the redundancy present in the dataset, we performed the analysis on a representative dataset that only consists of one structure per Uniprot accession number (Table 1) and found no difference in the clustering (S2 Fig.).

**Fig 3.**
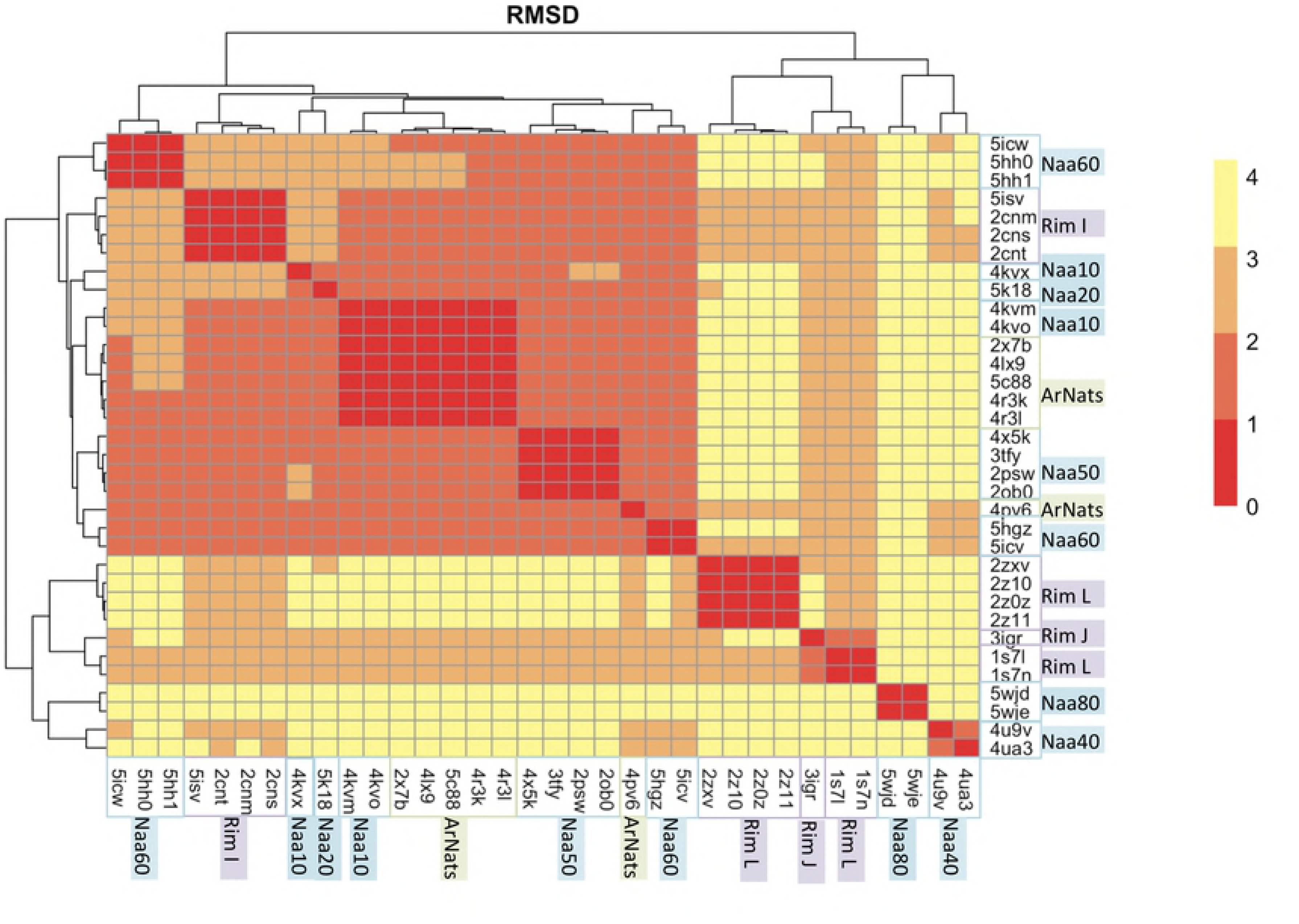
Heatmap representation of the pairwise Root Mean Square Deviations (RMSD). The dendrogram reflects the hierarchical clustering based on the RMSD values. The heatmap color scale goes from red (0Å < RMSD < 1Å; structural similarity) to yellow (3Å < RMSD < 4Å). Names of enzymes from eukaryotes are highlighted in blue, those of bacteria in purple and green is used for archaeal NATs.

The first group consists of: ArNats, Naa10, Naa20, Naa50, Naa60, RimI. In this cluster, the closest structures are the archaeal Naas and the eukaryotic Naa10 with RMSD values of 1.2 Å. The RMSDs between ArNats (e.g. 4l×9) and Naa10, Naa50, Naa60 and the RimI are lower than 2Å. The structural proximity of the archaeal Naa with enzymes belonging to other groups is in agreement with what is known about its substrate specificity. The archaeal Naa from *Sulfolobus* uses two different catalytic strategies; like Naa10 enzymes it can acetylate serines or it can acetylate methionines like the Naa50 enzymes (S2 Table). Mutations of key residues from the α1α2 loop were shown to shift the substrate specificity from small amino acids to methionines [42]. In this study, Liszczak et al. suggested these mutations as part of a model of the evolution of a eukaryotic ancestor to a more diverse family with different substrate specificity. The second main group consists of three clusters Naa40, Naa80 and the bacterial RimJ and RimL, which appear to be the most structurally distant from other structures in the dataset with RMSD values between 2.5 and 3.9 Å. They are composed of longer elements in the GNAT fold that influence the orientation of the secondary structure moving them further away from the other NATs. As shown in Fig. 2B the entire region from α1 to α2 is longer in RimL than in other Naas (6, 4 and 7 additional residues for helices α1, α2 and α1α2 loop, respectively). Naa40 also has an extended α1 helix of eight amino acids and an extra N-terminal helix α0 consisting of 17 amino acids. This α0 helix sits under the GNAT fold and changes the topology of the region β1-α2. The α1α2 loop and the longer α1 helix cover the active site and the β6β7 hairpin loop is flanked away from the active site (Fig. 2C). The structure of the β6-β7 region in Naa80 is different from that of typical NATs. It has a shorter β6-strand, which leads to a different orientation of the β6β7 loop and a ligand binding site opened more widely than in the other NATs [48].

### (2) Measure of pairwise flexibility similarity between NATs yield a grouping that coincides with similarity in substrate selectivity

The Bhattacharyya score (BC score) quantifies the intrinsic dynamics (dis)similarity between each pair of aligned cores of proteins in a dataset [54]. To this goal we perform normal mode analyses using elastic network models for each of the structures present in our dataset. We then calculated and compared the BC scores of the representative dataset (Table 1). A heatmap representation of those values together with a dendogram representing the clustering is shown in Fig. 4A. We observe that the structures are sorted in three main groups containing (1) Naa10, Naa20, Naa50, Naa60, archaeal NATs, bacterial RimI; (2) bacterial RimJ and RimL, as well as Naa40 which is the only eukaryotic NAT in this group and (3) Naa80. Interestingly, the first group is sorted between the Naas acetylating methionine: ArNats, Naa50 and Naa60 and the one acetylating small residues or alanine exclusively: Naa10 and RimI, respectively (S2 Table). Naa20 acetylates methionines followed by acidic residues and is clustered with Naa10. The latter has also been shown to shift substrate specificity towards acidic residues in its uncomplexed form [55]. Naa40 is also one of the most selective Naas since it acetylates only the Serine of the N-terminal of histones H4 and H2A. The bacterial Rims also have a narrow specificity and acetylate only ribosomal proteins (S2 Table). Naa80 is sharing the lowest BC scores with the other NATs. In addition of having a substrate-binding site wider than the other NATs, it also has a restricted substrate specificity towards the N-terminus of actin.

**Fig 4.**
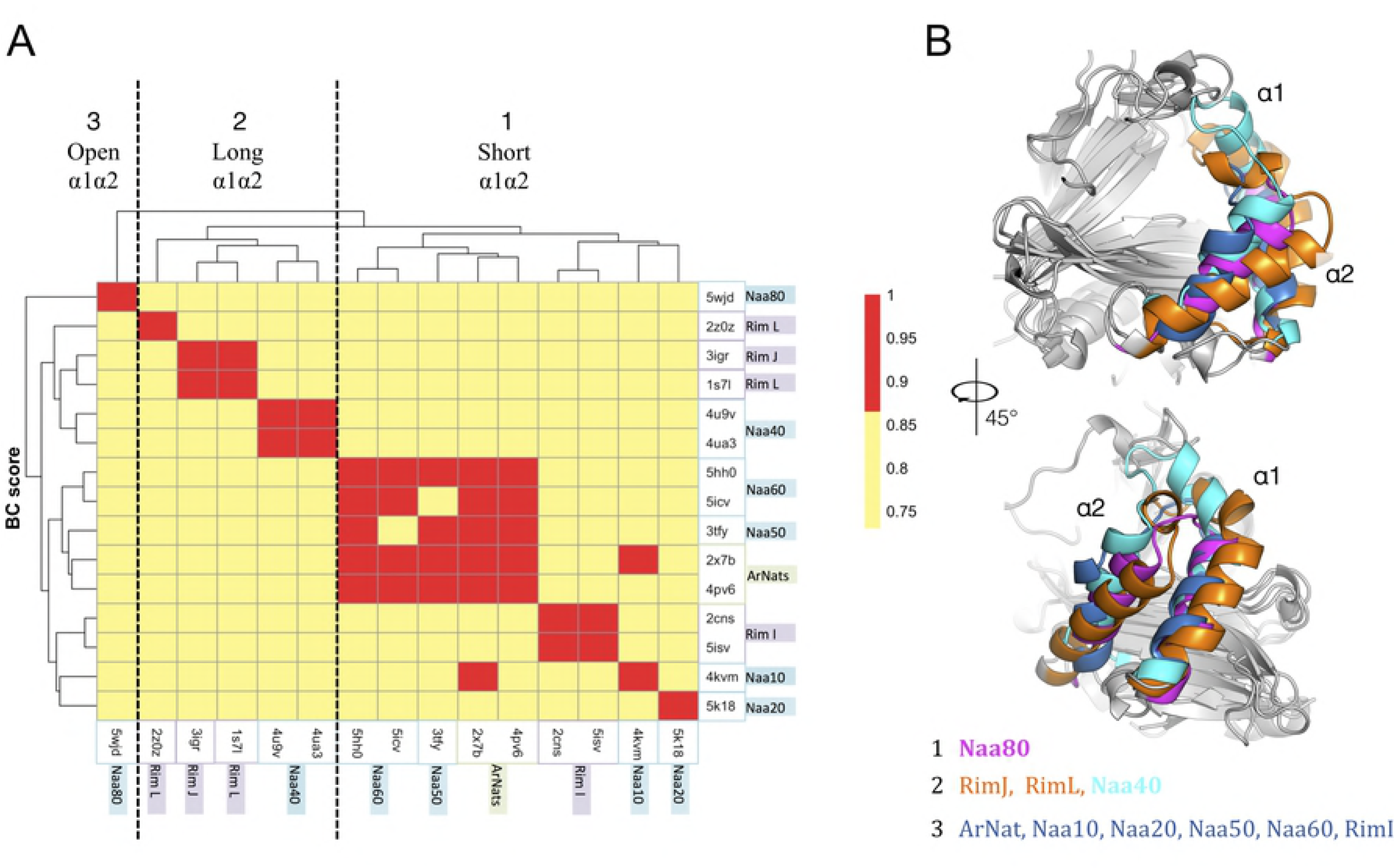
Comparison of the dynamics of the NATs using the Bhattacharyya score (BC score) on a non-redundant dataset. (A) Heatmap representation of the pairwise BC score between the representative structures (see Experimental procedures). The color scale of the BC score goes from red for high similarity in intrinsic dynamics to yellow for higher dissimilarity. The dendogram is the hierarchical clustering of the pairwise BC score. The names of the groups are written in boxes colored in blue for eukaryotes, purple for bacterial and green for archaeal NATs. (B) Cartoon representation of the structures aligned and used to calculate the pairwise BC score. The helices α1 and α2 are colored according to the cluster they belong to (see color of boxes on the axes of the dendrogram). The first cluster composed of the archaeal Naa, Naa10, Naa20, Naa50, the Naa60 and the RimI (colored in dark blue) shares a shorter helix α1 than that of the second cluster consisting of Naa40 (colored in cyan), RimJ and RimL (colored in orange). Naa80 (colored in magenta) separates from the others; it has the shortest α1α2 loop and the widest binding site of all NATs.

The BC-clusters show that flexibility (dis)similarities are encrypted in the coarse-description of the structure (actually restricted to its backbone) and in the harmonic potential of the ENM model (directly related to the proteins packing density).

### (3) ENM-NMA reveals the signature flexibility pattern of the GNAT fold

Similarity in fold or topology is generally associated with similarity in flexibility and dynamics [33,56]. We here intend to characterize the flexibility intrinsic to the topology of the GNAT fold. We then compare the results from ENM-NMA for each structure to reveal flexibility patterns intrinsic to the common fold. We first compute the normalised fluctuations for each amino acid in each structure, and the correlations between pairs of amino acids (Fig. 5A). The latter reveals how local motions are coupled across different regions of the fold.

**Fig 5.**
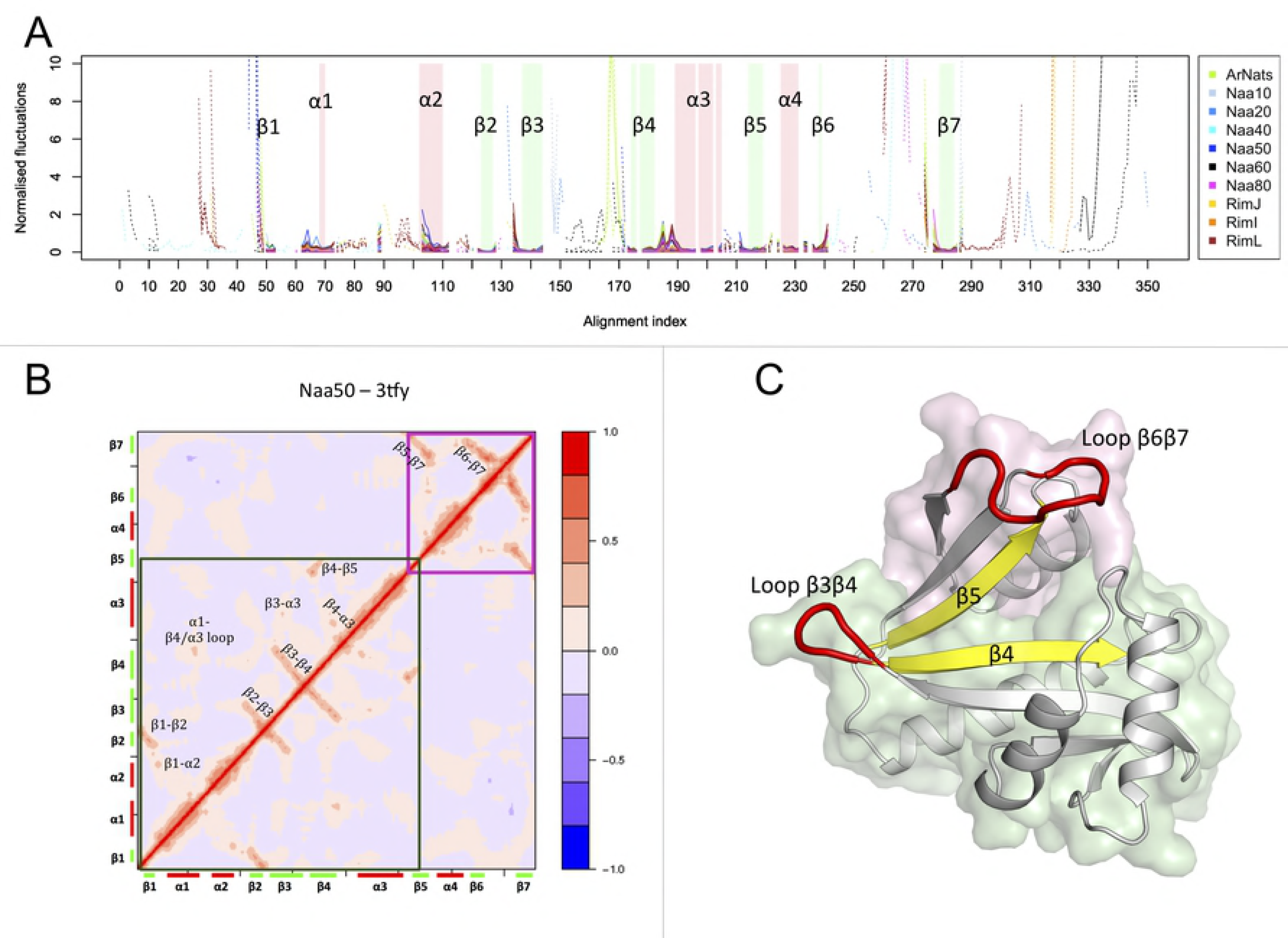
Normalized fluctuations and correlations. (A) Aligned normalized fluctuations for all NAT structures. For clarity one color is used for structures belong to the same group. The fluctuations depicted in plain lines are for positions where all structures align, while the dotted lines reflect the fluctuations of positions where not all NATs align. The secondary structures aligned in than 80% of all NATs are shown using green bars for the B-strands and red bars for the helices. (B) The correlation map of Naa50 (PDB ID: **3TFY**) shows the correlation patterns found in all maps (shown in Supporting Information Figure S3). Long-range correlations are found within two blocks highlighted within the green and pink frames. For all NATs the highest correlations are found within these blocks and not in-between. (C) Schematic representation of the main fluctuations and correlations on a cartoon representation of Naa50 (PDB ID: **3TFY**). The two blocks of correlations are distinguished by the color of their surface (green and pink). Regions with highest fluctuations are shown in red and those with lowest fluctuations in yellow.

#### Ac-CoA and the peptide substrate do not affect fluctuations significantly

X-ray structures in our dataset have been solved in different forms including with or without Ac-CoA and/or peptide substrate. For the sake of consistency, we should perform all computations using an ENM of the enzymes only. We thus first evaluate the effect of removing the Ac-CoA and peptide substrate from X-ray structure on the computed flexibility of the fold. We chose to compute the modes and normalized atomic fluctuations for the yeast Naa10 as crystal structures of different forms of the enzyme are available, notably two slightly different conformations, structures including Ac-CoA as well as structures including a bisubstrate inhibitor that mimics the presence of both Ac-CoA and peptide substrate [17]. Ac-CoA is included in the ENM as 11 beads chosen to represent an atom every 4 Å (see Experimental procedures) Normalized fluctuations are shown in S1 Fig. The positions of the minima and maxima are not affected by removing the Ac-CoA or substrate. When Ac-CoA or the bisubstrate are removed from the three structures (PDB IDs: **4KVO, 4KVX** and **4KVM**) the mobility of β4α3 and of the N-terminal end of helix α4 increases. Further, removing the bisubstrate from 4kvm also influences the fluctuations of loop β6β7 since it lies close to it. Generally, fluctuations are affected locally (i.e. at the Ac-CoA and ligand-binding sites) but not to a significant extent compared to the differences observed between NATs (Fig. 5A).

#### The V-shaped β-strands characteristic of the GNAT fold form a rigid core of the enzyme, while loops surrounding the binding site are mobile

The normalized fluctuations are plotted on Fig. 5A. We observe that the β-strands are the most rigid elements in all structures with β4 and β5 strands having the lowest fluctuations. Interestingly these two strands carry most of the catalytic amino acids (Fig. 2B) and our observation matches earlier reports of catalytic residues being positioned at particularly rigid points in protein structures [32,33,57–59]. Furthermore, β4 and β5 are not assembling into a sheet along their whole length despite their proximity. Instead they form a “V shape” splitting the seven-stranded beta-sheet and creating a crevice where Ac-CoA and peptide substrate meet (Fig. 1).

In the region of the helices α1 and α2, we notice a similar pattern of flexibility between all the structures where the loop α1α2 and the helix α2 fluctuate more. Molecular dynamics simulations of the human Naa50 and Naa10 have shown that the flexibility of helix α2 is decreased in the presence of a substrate [19]. This region is also involved in the complex formation with the subunit Naa15 [30]. We observed high fluctuations for long unfolded N- and C-terminal ends. Besides those regions where fluctuations cannot be calculated reliably, the highest fluctuations are observed for the β3β4 and β6β7 loops. Two tyrosines located at positions 239 and 240 of the alignment (Y138 and Y139 in Naa50), and conserved across several groups (not in RimL, Naa40 and Naa80), are located on loop β6β7. They are known to be involved in substrate binding and form hydrogen bonds with the substrate [19].

#### The GNAT fold is divided in two dynamical domains on either sides of the β5 strand

We calculated the correlations for each of the representative structures. The representative correlation map is shown as a heatmap in Fig. 5B. All NATs share the same pattern consisting of two blocks with relatively little correlations between them (S3 Fig.) indicating that the proteins contains two dynamical domains [60]. The boundary between the two coincides with the V-shape split between β4 and β5; the first domain starts at sheet β1 and ends before strands β5, and the second domain starts at strand β5 and ends with strand β7. Within each domain, pairs of neighbouring β-strands are strongly correlated, as expected for beta-strands involved in the same sheet [61], but to a lesser extent for β4-β5. This is explained by the distance between β4 and β5 and the split in the fold at the β4-β5 interface. Correlations between strands and helices are weaker in general and happen through extremities of helices only. This is in agreement with what we observed for enzymes with the TIM barrel fold [33]. β5 shows strong correlations with strands on either side; β4 and β7. Correlations between domains are weak in comparison. There is an anti-correlation of such type between loops α1α2 and β6β7, which are in contact with each other forming a lid above the catalytic site. The potential flexibility of the loop β6β7 to open and move independently of the other block reinforces the idea that it could create more room for the substrate to enter. Apart from the loop, it seems that the movement of the pink block (Fig. 5C) independently from the first block will influence the size of the binding pocket.

### (4) Structural differences in the N-terminal-α2 region contribute to difference in dynamics between NATs

Differences in intrinsic dynamics are encoded in the structure. The GNAT fold has a region of high variability from the N-terminal to the α2 helix [7] (Fig. 2C and 4B). As noted earlier the helices α1 and α2 are longer in Naa40, RimJ and RimL than in the other NATs from group 1. RimJ and L have 1.5 additional turns in each of the two helices compared to structures clustered in group 1, while the two extra turns of helix α1 in Naa40 brings its C-terminal end over the active site at a location overlapping with that of the β6β7 hairpin loop in the other NATs. As a result β6β7 is protruding further away from the protein core than in the other NATs and shows very large fluctuations as calculated from the modes (Fig. 2C and Fig. 5A). Furthermore Naa40 has an extra N-terminal helix α0, the movements of which are correlated with strands β3 and β4 as well as with loop α3β5 (Fig. 6). On the contrary Naa80 has a shorter α1-α2 loop as well as a different orientation of its β6β7 loop.

**Fig 6.**
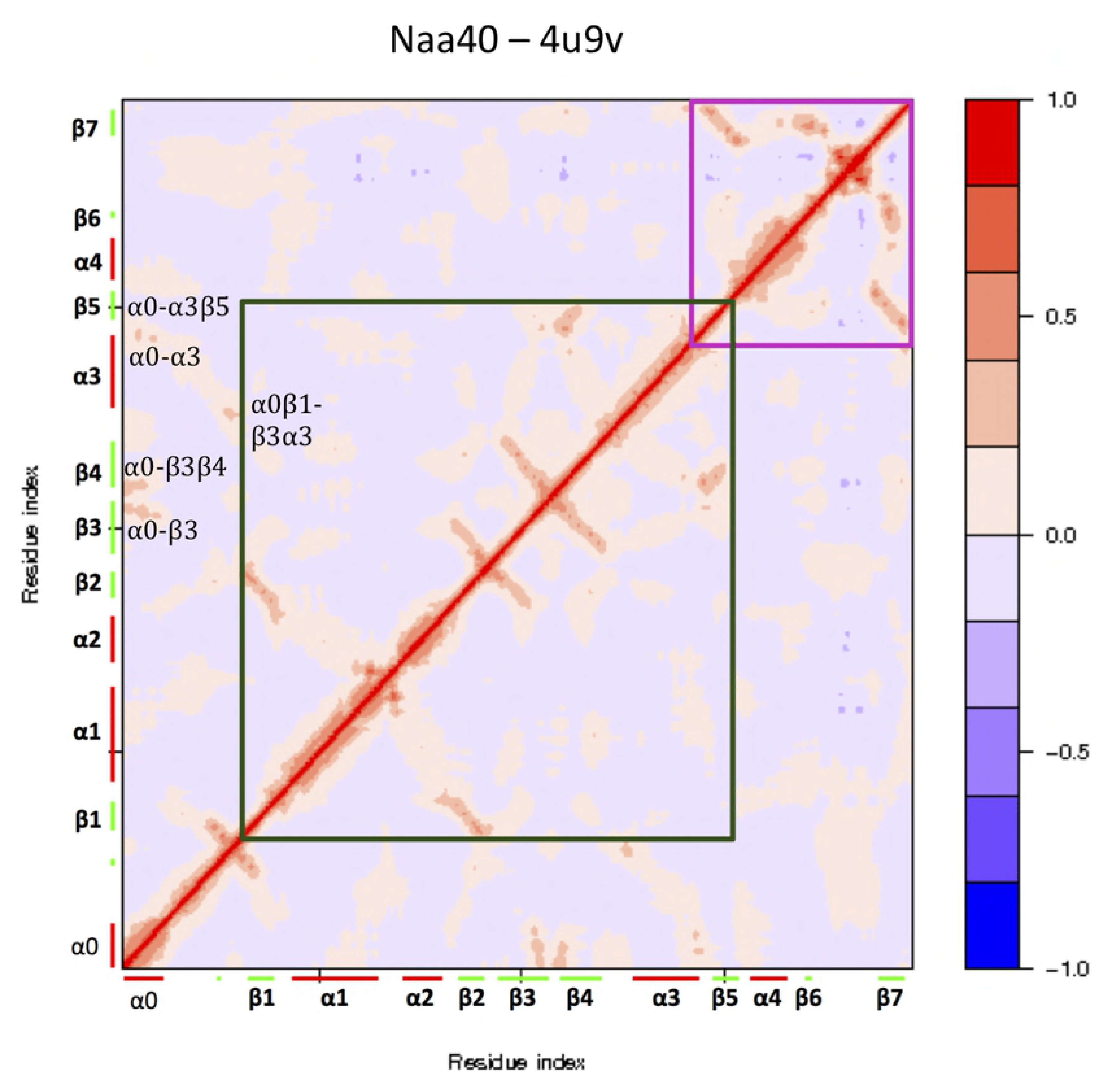
Correlations maps for the representative of the Naa40 group. Helix α0, which is not present in other NATs, has high correlations with sheets β3 and β4, and with the α3β5 loop.

The fact that clustering structures according to the pairwise BC score sorts them into groups that differ by their N-terminal end to α2 region indicates that the dynamics of this region is significantly different between the three groups.

### (5) The mobility of the β6β7 loop influences peptide substrate accessibility to the catalytic site

The low frequency modes yield deformations at the lowest energetic cost and have been shown to be functionally relevant [62,63]. Kurkcuoglu et al. showed that global motions of functional loops were important for fitting and binding of the substrate [64]. Given the high mobility of the β6β7 loop and its position with respect to Ac-CoA and the substrate binding site, it is legitimate to wonder how its movements would influence the fairly narrow tunnel in which substrate and Ac-CoA meet. In order to determine the contributions of loop motions to the accessibility of the active site, we evaluated the changes to the surface area of the tunnel entrance when the structure conformation is modified following the lowest energy normal modes.

Further, we calculated how the surface area changes when the structure is modified following the individual lowest energy normal modes (see Experimental procedures). A reconstitution of the protein side chains from our C-alpha model was necessary to calculate the surface area of the mouth of the tunnel using CAVER Analyst [65] (see Experimental procedures). To this goal we first defined a static clipping plane using three residues (Tyr31, 73 and 138) lining the entrance (see Experimental procedures). We generated a trajectory along the first six lowest-frequency normal modes and held this static clipping plane for each frame of the trajectory. Note that the orientation of the clipping plane was extracted from the conformation of the X-ray structure and the same orientation was used in each frame. We then calculated the value of the surface area throughout the trajectory. We also calculated the surface area at a clipping plane 2Å away from the *mouth*, where the second and third residues of the substrate sit, in order to verify that no amino acid closes the access upwards of the active site.

The lowest energy modes show concerted motions of the three loops β3β4, β6β7 and α1α2. They surround the active site and their movements modify the shape of its *mouth* and its surface area (Fig. 7). However, the motions of each loop seem to have a different effect on the surface area. As illustrated in Fig. 7B slight movements of Tyr138 in Naa50, found on the β6β7 loop, drastically change the opening of the entrance (from 50.5Å to 71.1Å). The β6β7 loop provides a simple steric regulation of the entrance of the binding site due to its position as a lid above the active site.

**Fig 7.**
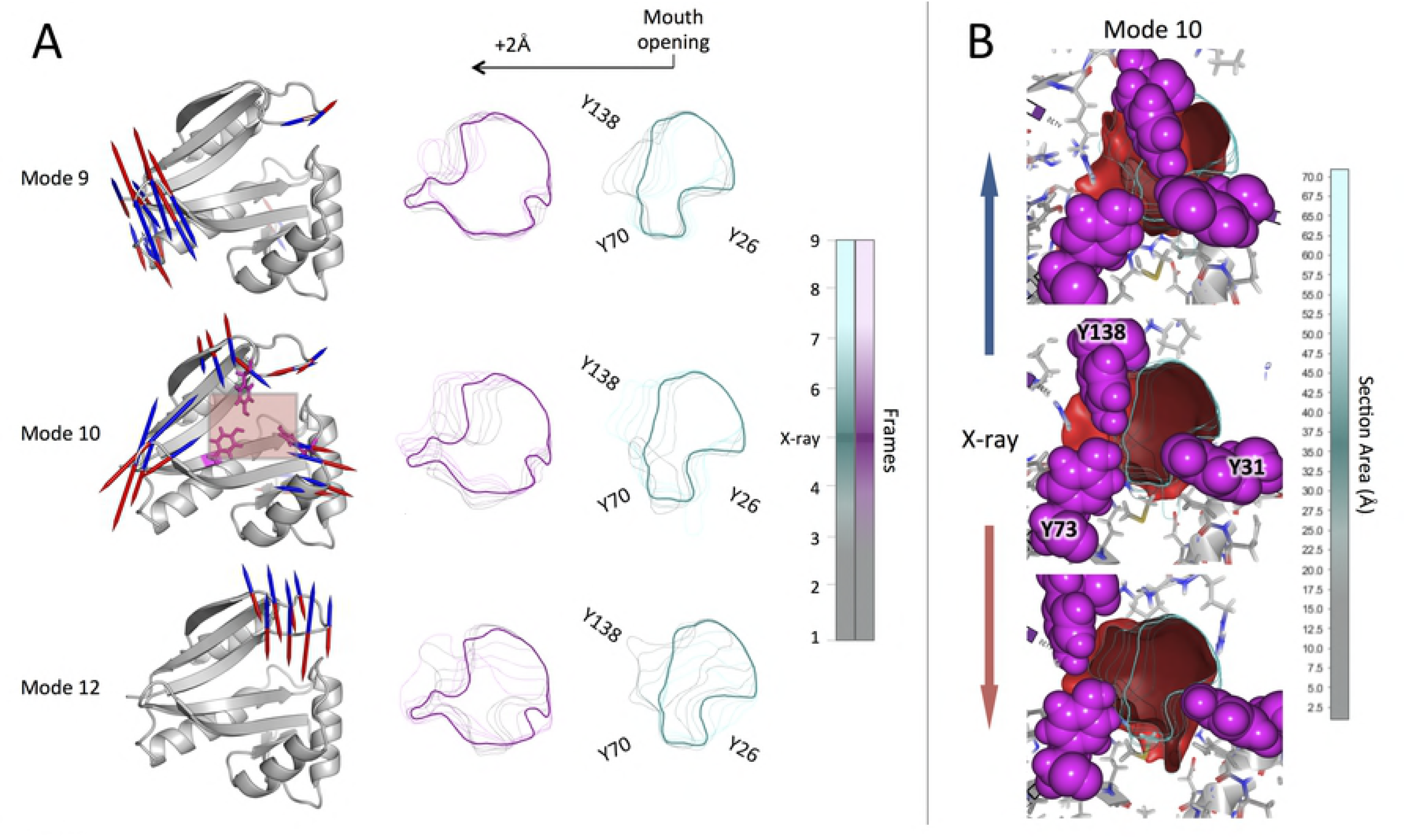
Influence of the flexibility of loop β6β7 on the surface area of the cavity mouth. Amino acids Y138, Y70 and Y26 line the *mouth* of the cavity in the X-ray structure of Naa50 (PDB ID: **3TFY**). (A) Representation of the low-energy modes affecting the surface area of the *mouth* the most. Naa50 is in cartoon representation with arrows indicating the direction of the movements corresponding. Only the vectors with a minimum length of 4Å are represented, with in red colour the positive factors and in blue the negative ones. The trajectory is 9 frames long, with the X-ray structure coordinates found in the middle (frame 5). Four frames are used to represent each direction of the vectors (from frame 1 to 4 for the positive vectors and from frame 6 to 9, for the negative vectors). The contours in cyan shades represent the evolution of the mouth surface area throughout the trajectory. The bold contour is obtained from the X-ray structure. The surface areas are also calculated two angstroms before the mouth of the cavity (contour in pink shades) to ensure that no amino acid comes closing the mouth. (B) Close up of the red square found on the cartoon representation of the mode 10 in (A). Clip plane of the protein at the mouth area defined by the three tyrosines (shown in pink spheres). The red surface represents the rest of the tunnel calculated by CAVER Analyst where the N-terminal and the Ac-COA meets (see Experimental procedures). The contours are the surface areas of the mouth depicted in cyan shades in (A). The scale indicates the range change in size of the surface throughout the trajectory. The bold contour is the contour in the X-ray structure and has a size of 50,5Å. Small movements of the Y138 in the low-energy mode 10 impact greatly the surface area of the entrance of the binding site (from 2.5 to 70Å).

## Discussion

The variety of NATs enables the acetylation of a wide variety of N-termini of proteins at different localizations in the cell. The GNAT fold shared among all NATs and with other acetyltransferases such as KATs offers a common scaffold to perform the catalytic activity. One particular feature of the GNAT fold in the NATs is a tight entrance of the binding site, where the β6β7 hairpin loop shields the active site and forms a tunnel together with the α1-α2 region (except in Naa80). The loop seems to enable the entrance of only N-terminal amino acids (Fig. 1C). However, it remains unclear how substrates are able to enter through the tight *mouth* of the tunnel. Furthermore the path through which bisubstrate inhibitors bind remains uncertain. Using ENM-NMA we have compared the intrinsic dynamics of the GNAT fold in all available structures of NATs covering all domains of Life. We have shown that i) there is a dynamic pattern in the fold common to all NATs, ii) the β6β7 loop is highly mobile and iii) intrinsic dynamics of the loop regulates the accessibility of the binding site.

We identified a common pattern in the intrinsic dynamics of the GNAT fold, where the β-sheet core is more rigid than the rest of the structure. The two strands, β4 and β5, are the least flexible elements of the β-sheets. Interestingly, the β4 and β5 strands carry residues involved in the proton wire essential for the transfer of the acetyl. The rigidity of the catalytic core in the GNAT fold is in agreement with case-studies of enzyme dynamics where residues involved in catalysis are found to be placed at rigid conserved positions of the fold thus stabilizing amino acids essential for the function [33,56,58].

The catalytic core is at the crossroads of two dynamic domains that move independently from each other. The region between the C-terminal end of α3 and the N-terminal end of β5 is found to have hinge residues that are important pivots in larger motions of the protein. The pathological Naa10 p.V107F mutation at the N-terminal end of β5 causes a 95% reduction of the catalytic activity compared to Naa10 WT [66]. In this study, Popp et al. built a homology model of the mutant V107F and observed a disruption of hydrophobic contacts with the Met98 found on helix α3 of the human Naa10. This correlates with our results and shows that perturbation of the packing density of hinges region affect function through a modification of the intrinsic structural flexibility [67].

The most mobile regions in NATs are the functional loops α1α2, β3β4 and β6β7. Substrate-binding residues have a tendency to be in more flexible regions than catalytic residues [56,68,69] and the α1α2 and β6β7 loops carry residues involved in substrate binding [19]. The C-terminal end of helix α2 has been identified as carrying a mutation (p. S37P) in the human Naa10 that is causative of the lethal Ogden syndrome [4]. Using molecular dynamics simulations on the model of the human NatA complex, we have earlier shown that this mutation decreased the fluctuations of the α1α2 loop and of the α1 helix. Moreover the mutation impairs the catalytic activity and the formation of the NatA complex, inducing a reduction of NatA-mediated N-terminal acetylation and affecting cell proliferation [4]. The α1 and α2 helices are also part of the binding interface with Naa15, the auxiliary subunit of the NatA complex [17] (S2 Table). The loop β3β4 is 10 to 15 residues longer in Naa60 than in the other NATs and mutations on key residues disrupt interactions with the β5, β6, and β7 strands, leading to altered catalytic efficiency and protein stability [70]. Finally the β6β7 loop, one of the most flexible loops, contains two tyrosines conserved through most of the NATs, which make hydrogen bonds with the backbone of the first and second amino acids of the substrate [19].

The highly mobile α1α2 and β6β7 loops flank the catalytic core regulating the size of the binding site. Moreover the regions from the N-terminal to α2 and β6-β7 have structural differences among the NATs. The BC score, which quantifies similarities of intrinsic dynamics between the aligned core regions of proteins, highlighted dissimilarities between Naa40, the bacterial RimL and RimJ in one hand, and the rest of the NATs in the other hand; while Naa80 clusters on its own. Interestingly, Naa40, RimL and RimJ have longer α1 and α2 helices and a longer α1α2 loop, which results in slight changes of the active site shape. As far as we are aware of, Naa40 is the only known NAT with a different position of the substrate in the active site. As shown in Magin et al., all the substrates in other structures have their 2^nd^ and 3^rd^ residues sitting close to the α1 and α2 helices, while the α2 helix in Naa40 obstructs this region shifting the substrate towards the β5 and β7 strands [47]. The bacterial RimL is only active as a homodimer unlike the other NATs, which tend to be active as monomers complexed with auxiliary subunits forming heterodimers or heterotrimers (S2 Table). The β6β7 loop is part of the RimL dimerization interface. Helix α2 is tilted away from β4 yielding a larger mouth of the cavity compared to the other NATs. The longer elements in Naa40, RimJ and RimL thus illustrate how secondary structure elements lining the binding site affect its size, shape, and accessibility. In the case of Naa80, the opening to the binding site is wider than in other NATs and this is thought to play a role in its specificity for the acidic actin N-termini [48].

The collective motions known to support function have been shown to be well described by the lowest energy modes [63,71]. Those modes in NATs contain large movements of the β6β7 hairpin loop with hinges found at the extremities of the neighbouring strands. We have demonstrated the impact of this motion on the shape and size of the entrance of the substrate binding site using Naa50. It acetylates methionine residues followed by hydrophobic residues [72,73]. As shown in Fig. 1B, the entrance of the active site tightly surrounds its substrate. From the X-ray structure we can measure that the methionine is approximately 6Å long and observe that Val29, Tyr31, Tyr138 and Tyr139 tightly surround it. The movements of Tyr138 alone, which is located on the β6β7 loop, cause a widening of the mouth in the low energy modes. This tyrosine is actually not present in Naa40, and the bacterial RimL where it is replaced by a glycine in Naa40 and asparagine or alanine in RimL. These changes might affect the substrate specificity. In general it has been observed that regions playing a role in ligand-binding have a tendency to explore more of the deformation pattern available than the catalytic site without impairing enzyme function [74].

Taken together, we propose that the dynamics of the two domains and the high mobility of the β6β7 loop give the ligand binding site a flexibility that is important for its selectivity. For the first time, we suggest a possible conformational change that would explain how the long bisubstrates are able to bind both the Ac-CoA and peptide substrate binding sites. Our calculations show that hinges of these movements are located at V131 and A145 in Naa10 (PDB ID: **4KVM**). Further, this loop represents a striking structural difference between NATs and KATs and is hypothesized to prevent the entrance of internal lysines in the active site [15,47,75]. Our calculations show that fairly small displacements of the β6β7 loop as a rigid entity modify the accessibility to the active site and the Ac-CoA. β-hairpin loops are known to have a strong hydrophobic core and interstrand interactions keeping them compact and the turn stable [76]. The human Naa10 has been shown to acetylate internal lysines of various proteins [24,26,27,77] and the auto-acetylation on its K136 found on the β6β7 loop could be the reason of its shift of substrate specificity towards internal lysine [78]. Acetylation of unexpected substrates might be enabled by an increased mobility of loop β6β7 triggered by either a particular substrate or experimental conditions.

The importance of the dynamics-function relationship of the GNAT fold in the NATs is undeniable. Our work fills a gap in the understanding of the versatility and broad substrate specificity of the NATs enzymes [4,19]. Our results are relevant for those seeking to design inhibitors of NATs involved in cancer, Huntington’s disease or other pathologies. In addition, we reveal the role played by the flexibility of the β6β7 hairpin loop in the opening of the ligand binding site. Further investigations are needed to experimentally evaluate the extent of the influence of the loop mobility on NATs activity and substrate specificity. This could be done by mutagenesis experiments where selected amino acids in hinge regions could be replaced by glycine or proline to increase or reduce loop mobility. Such an approach would present the advantage of not affecting the structure and stability of the β-hairpin itself [76,79].

## Materials and Methods

### Dataset preparation

Scrutinizing protein structure and fold databases such as CATH or SCOP we could not find a specific grouping for all NATs. We used the annotation of GNAT domain from PROSITE [80] (PROSITE code: PS51186) and “N-terminal protein amino acid acetylation” as a biological process in PDBe [81]. We collected more than 160 structures and filtered down to 45 structures that had annotations as N-alpha acetyltransferases or N-terminal acetyltransferases (Table 1). From these we excluded eleven structures for which the X-ray structure had unresolved segments within the GNAT fold (S1 Table).

We formed 10 functional groups: Naa10, Naa20, Naa40, Naa50, Naa60, Naa80, archaeal NATs, RimI, RimJ and RimL. These groups have been constituted either by considering the Enzyme Commission (EC) number, their functional annotation in scientific literature when available or the kingdom of the organism the protein is found in. All structures files were prepared for the calculations by selecting one chain in the assembly and removing the Ac-CoA or peptide substrate if present. The reference set consists in one structure, called representative, for each Uniprot code in each functional group (see PDB IDs in bold in Table 1).

### Structural alignment

The structural alignment was obtained using MUSTANG [51] which has been shown to be efficient on distant related proteins [33,34]. The algorithm performs a progressive pairwise alignment using the position of Cα atoms. It extends the pairwise structural alignments into multiple structure alignments by recalculating a pairwise residue-residue score at each step of the extension and progresses using a guide tree. The pairwise RMSD between Naas structures cluster is displayed as a heatmap plotted with the R function pheatmap [82]. The alignment provided is primarily used for the comparison of the intrinsic dynamics as described in Refs. [33,83].

### Elastic network model and Normal mode analysis (ENM-NMA)

The normal mode analysis has been performed using WEBnm@ [83]. The web-tool is using an Elastic Network Model (ENM) modelling protein structures as a network of nodes, the Cα atoms, connected together by Hookean springs. We used the Calpha force field [84,85], as implemented in the Molecular Modelling Toolkit [86]. It uses a pair potential to describe the interactions between two Cα atoms as:

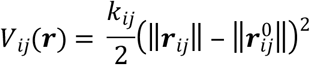

where *r*_*ij*_ is the distance vector between two Cα atoms *i* and *j* in the configuration *r* of the protein, 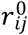 is then the same pairs of atoms *i* and *j* at the equilibrium conformation and *k*_*ij*_ is the non-uniform force constant defined by:

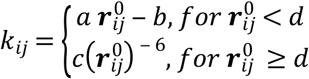

with *a* = 8.6 × 10^5^kJ mol^−1^nm^−3^; *b* = 2.39 × 10^5^kJ mol^−1^nm^2^; *c* = 128 kJ mol^−1^nm^4^ and *d* = 0.4 nm.

The potential energy of the network model is then the sum of all the atomic configurations:

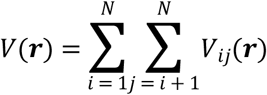

where *N* is the number of nodes in the network.

For the normal modes calculation of the holo form of *Schizosaccharomyces pombe* Naa10 we represented CoA by 11 beads placed at the positions of atoms distant of 3 to 4Å, namely C, C3P, C6P, C9P, CCP, P1A, C4B, P3B, N9A, N6A and N3A.

### Normalized fluctuations

The fluctuations *F*_*i*_ give the variances of each atom position and are given by:

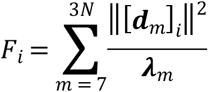

where ***d***_*m*_ is the displacement vector of the atom *i* in mode *m. F*_*i*_ is then the sum of the all the displacements of *i* for all the non-trivial modes that are weighted by their eigenvalues.

### Correlations

The matrix of correlations is calculated from the normal modes [87] which quantifies the coupling between two atoms *i* and *j* as:

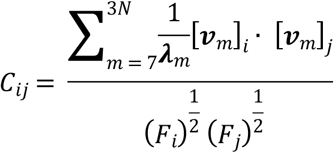

*C_ij_* = 1 when the motions are completely correlated and *C*_*ij*_ = −1 when they are completely anti-correlated.

### Bhattacharyya coefficient score

Finally we used the Bhattacharyya coefficient (BC) score to compare the effective covariances of the common aligned cores of two structures A and B as implemented in WEBnm@ and is based on Fuglebakk et al. [54] :

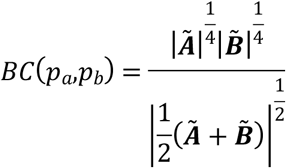

### Trajectories and side chains reconstruction

We selected the 6 first non-trivial modes from the set of normal modes of the protein of interest. For each mode, we generated a trajectory consisting of nine snapshots; we displaced the initial Cα positions following the mode and using arbitrary amplitudes in either direction around the X-ray structure (the mode vectors were multiplied by −12, −9, −6, −3, and 3, 6, 9, and 12).

We used the Molecular Modelling ToolKit (MMTK,[86]) to reconstruct the side chains in order to obtain trajectories at all-atom resolution. We calculated the 3D transformation necessary to superimpose the initial all-atom structure onto each of the snapshots of the Cα trajectories. This was done by minimizing the RMS difference between the initial all-atom structure and the Cα trace snapshots. The 3D transformations were not computed on the overall structure but locally using an iterative process. We used sliding windows that were three amino acids long to compute the transformation, which is then applied only to the central amino acid for which the side chain is reconstructed. The process is then iterated by sliding by one residue along the protein sequence.

### Visualisation and calculation of the cavity

The analysis was performed using CAVER Analyst [65]. We first selected three Cα atoms from amino acids (Tyr31, 73, and 138) that are lining the mouth of the cavity in the X-ray structure. In 3D space, these three atoms define a parallel plane to the tunnel mouth. Then a set of intersecting spheres, with a radius of 1Å, is placed on a line perpendicular to this plane to fill up the length of 5Å into the cavity. Using this geometrical structure as a base, we computed a cavity surface in each frame using the algorithm described in [65]. We do not use the extension of the algorithm proposed by Jurcik et al. [88] as it was developed for the detection of deeply buried voids inside proteins. While in our case the cavities in NATs are not fully surrounded by amino acids and are closer to the protein surface. Finally, the contours of the surface area of the mouth are extracted using the MoleCollar algorithm part of CAVER Analyst and first described in Ref. [89]. The MoleCollar algorithm estimates the empty space area by triangulating a contour that is obtained by cutting the cavity surface by a plane and summing up the areas of all resulting triangles. The algorithm was modified to keep the same orientation of the cutting plane throughout the trajectory in order to follow the evolution of the surface area along the trajectories.

## Acknowledgements

We would like to thank Konrad Hinsen (Centre de Biophysique Moléculaire in Orléans, France) for fruitful discussions on trajectory creation; Edvin Fuglebakk (Institute of Marine Research, Bergen, Norway), Sandhya Tiwari (Computational Structural Biology Research Unit, RIKEN, Japan), Bojan Krtenic (Department of Biological Sciences, University of Bergen) and Simon Mitternacht (University of Bergen Library) for their valuable comments on the manuscript.

## Author contributions

Conceived and designed the experiments: AA, NR, JB, PB and TA. Performed the experiments: AA. Analysed the data: AA, NR. Code development for CAVER Analyst: JB. Code development to recreate all-atoms trajectories: PB. AA and NR wrote the paper with contributions from all authors.

## Supporting Information

**S1 Fig. Effect of the presence of a cofactor or a bisubstrate on the normalized fluctuations of the Naa10 from *Saccharomyces pombe***. Naa10 from *S. pombe* has been crystallized with Naa15 (PDB id: 4kvm and 4kvo) and without Naa15 (PDB id: 4kvx). The main difference between the structures of the two states is a rearrangement of helix α2 and of the α1α2 loop. The experimental structures of the complexed Naa10 form contain either a bisubstrate inhibitor (4kvm) or only a cofactor (4kvo). The structure of the uncomplexed Naa10 form has only the cofactor (4kvx). The computations were performed on the X-ray structures with their ligand bound, and on the same protein structures after removal of the ligand.

**S2 Fig. Heatmap representation of the pairwise Root Mean Square Deviations (RMSD) for representatives structure.** The dendrogram reflects the hierarchical clustering based on the RMSD values. The heatmap color scale goes from red (0Å < RMSD < 1Å; structural similarity) to yellow (3Å < RMSD < 4Å). Names of enzymes from eukaryotes are highlighted in blue, those of bacteria in purple and green is used for archaeal NATs.

**S2 Fig. Correlations maps for the representative of each NAT group.** Dark red and blue show correlated motions between pairs of residues (close to 1 or −1, respectively), when lighter colours refer to correlations close to 0. Long-range correlations are found within two blocks highlighted by the green and pink frames. For all the NATs the highest correlations are found within these blocks and not in-between.

**S1 Table.** Structure dataset.

**S2 Table.** Ligand specificity of NAT enzymes included in the dataset. The information in this table is collected from the following references: ArNat [14,42], NatA [1,17,55,90], NatB [91–93], NatD [47,94], NatE [73,95,96], NatF [52,97], NatH [11,48], RimI [12,29], RimJ [12,98], RimL [13,49].

